## Supplementary material for "Reduced discrimination between signals of danger and safety but not overgeneralization is linked to exposure to childhood adversity in healthy adults"

### Supplementary Methods and results

#### Additional information on the paradigm.

In brief, two female faces with neutral expressions served as the CS+ and CS- (see also Supplementary Figure 3); CS duration: 6s, assignment randomized) and a loud female scream (95dB, delivered via over-ear headphones, US duration: 1.5s) co-occurring with the CS+ face showing a fearful expression that served as the US. Both faces were presented four times each during habituation and 12 times each during acquisition training. During the generalization phase, both faces were shown 12 times in addition to 12 presentations of each of four generalization stimuli (GS) that were morphs of the CS+ and the CS- faces in steps of 20%. During intertrial intervals (ITIs), a white fixation cross on black background was shown for 9-12s. The generalization phase consisted of two blocks (6 trials each). Since a discrimination training (Herzog et al., 2021) was conducted between blocks for some participants, only the first block of the generalization phase was analyzed here. The reinforcement rates were 83.3%, 50% and 0% during acquisition training, generalization (i.e., to prevent extinction), and habituation, respectively. Participants were not instructed about the CS-US contingencies and were merely informed they should passively view pictures. The participants were provided with an extinction session at the end of the experiment to guarantee that no lasting conditioning would take place (Schiele, Reinhard, et al., 2016).

#### Additional information on the study sample.

Participants of this study were recruited in a multi-centric collaborative research center ‘Fear, anxiety, anxiety disorders’ joining forces between the Universities of Hamburg, Würzburg, and Münster, Germany (SFB TRR58). During the second funding period (2013-2016), all three sites recruited a large sample (N ~500) in the context of the Z project. All participants underwent the cross-sectional experimental paradigm reported here and were additionally extensively characterized to allow specific subprojects to recruit target sub-populations serving different aims with a focus on molecular genetic, epigenetic, or other research questions (see Herzog et al. (2021); Imholze et al. (2023); Schiele, Reinhard, et al. (2016); Schiele, Ziegler, et al. (2016); Stegmann et al. (2019)). The question on the association of exposure to childhood adversity and recent adversity was part of the primary research question of one subproject led by the senior author of this work (B07, TBL) and was hence a research question of primary interest also for this multicentric project.

Additional exclusion criteria included left-handedness, non-Caucasian descent (as sub-projects focused on genetic analyses, e.g. Schiele, Ziegler, et al., 2016), pregnancy, severe medical diseases, intake of illegal drugs, psychoactive medication or excessive consumption of alcohol, nicotine, and caffeine (for details see Schiele, Reinhard, et al., 2016; Schiele et al., 2020).

Socioeconomic status (SES) in this sample might be roughly inferred from education level and current occupation status. Significant differences were observed between individuals who were exposed as compared to unexposed to childhood adversity in relation to their school degree ($\chi^{2}$(6) = 15.89, *p* = .014). More precisely, significantly less individuals (*p* = 0.006) exposed to childhood adversity as compared to those who were unexposed hold a high school diploma (in Germany: ‘Abitur’) with (still) unfinished studies. However, no significant differences were observed in terms of employment ($\chi^{2}$(8) = 14.60, *p* = .067) or type of occupation. ($\chi^{2}$(10) = 13.73, *p* = .185; see Supplementary Figure 9).

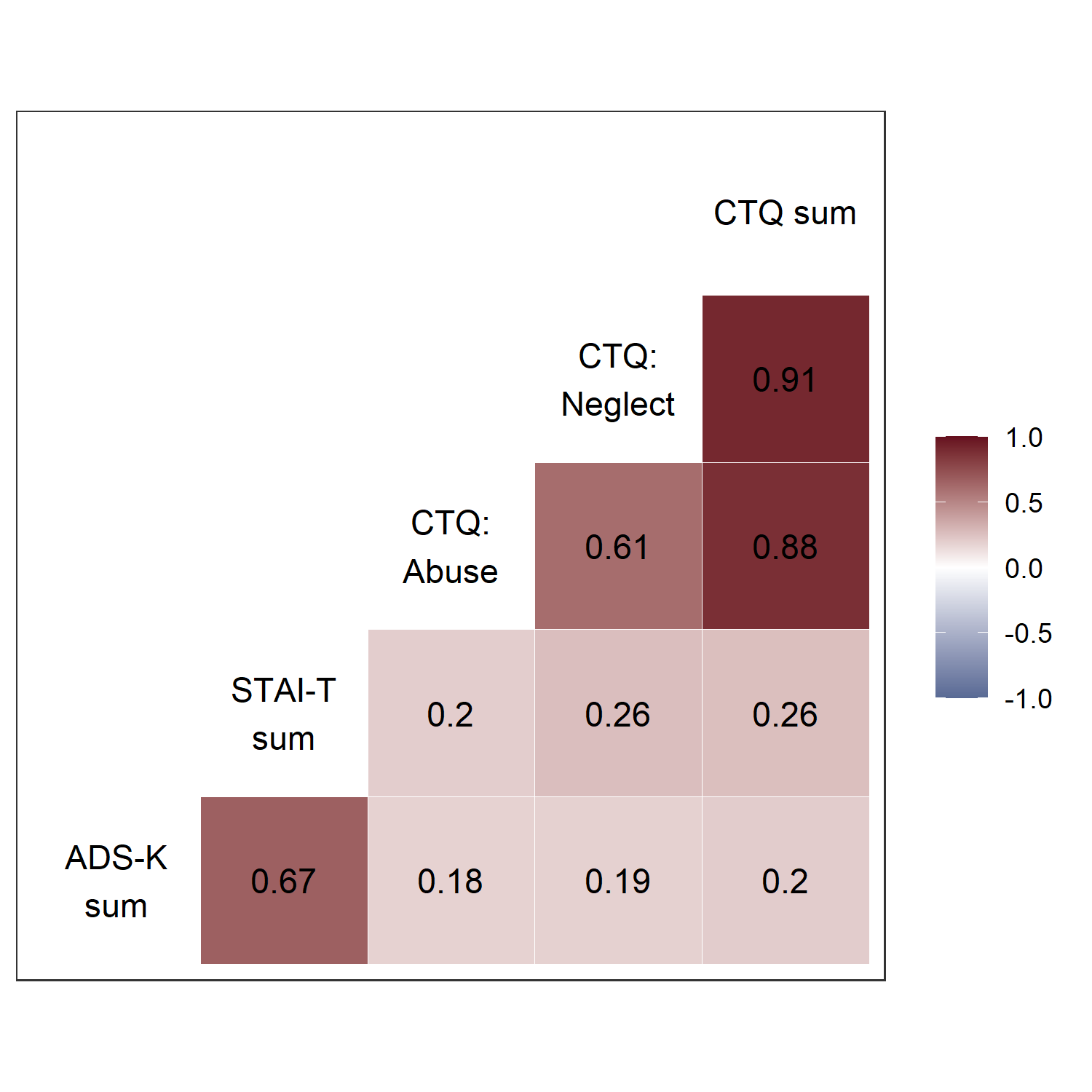

Supplementary Figure 1: Illustration of zero-order correlations (Pearson’s correlation coefficient) between relevant sample characteristics: different scores of childhood adversity (CTQ sum, CTQ: Neglect, CTQ: Abuse), trait anxiety (STAI-T sum) and depression (ADS-K sum). All depicted correlations were significant (all *p*’s < 0.001). ADS-K = short version of the Center for Epidemiological Studies-Depression Scale (Allgemeine Depressions-Skala, Hautzinger & Bailer, 1993), STAI-T = State-Trait Anxiety Inventory, Trait (Spielberger, 1983), Abuse and Neglect = composite scores built from childhood trauma questionnaire subscales (CTQ-SF, Bernstein et al., 2003; Wingenfeld et al., 2010).

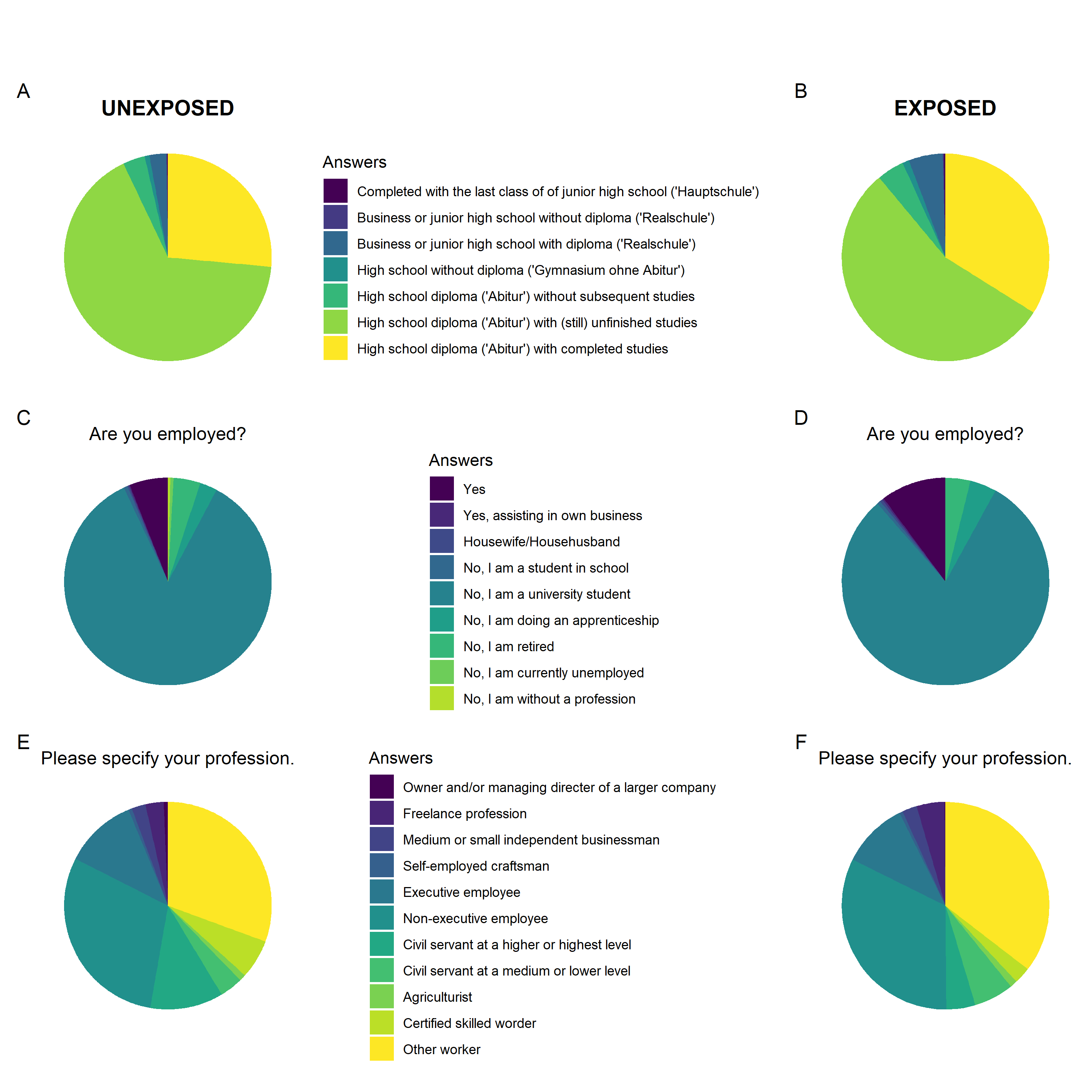

Supplementary Figure 2: Illustration of the socioeconomic status information of individuals unexposed (left) and exposed (right) to childhood adversity (according to an at least moderate childhood adversity cut-off) inferred from questions about the school degree (A and B), and the current occupational status (C and D) including the type of employment (E and F).

[Figure not shown:
„Our policy is to avoid the inclusion of photographs and any other identifying information of people, whether it be patients, participants, test volunteers, experimental stimuli or previously published photographs, because verification of their consent is incompatible with the rapid and automated nature of preprint posting” (bioRxiv, 2024)]

Supplementary Figure 3: This illustration of the study design was created as part of the publication ‘Prediction of Changes in Negative Affect During the COVID-19 Pandemic by Experimental Fear Conditioning and Generalization Measures’ by Imholze et al. (2023) (DOI: <https://doi.org/10.1027/2151-2604/a000523>), which was distributed as a Hogrefe OpenMind article under the license CC BY 4.0 (<https://creativecommons.org/licenses/by/4.0>). Please note that the face stimuli used in the actual experiment are different from those shown here. For copyright reasons, we are unable to present the blonde women and have used an image of a redheaded woman as a substitute.

#### Supplementary Analyses on measurement reliability.

Here, we follow recent calls for an increased focus on measurement reliability (Cooper, Dunsmoor, et al., 2022; Klingelhöfer-Jens et al., 2022; Zuo, Xu, & Milham, 2019) and report split-half reliability for SCRs for the fear acquisition training and generalization phase. Split-half reliability was calculated by correlating (Pearson’s correlation coefficient) averaged odd and even SCR trials (i.e., odd-even approach) for the acquisition and generalization phases separately (see Supplementary Table 1). For acquisition training, the first trial was excluded (Klingelhöfer-Jens et al., 2022).

The within-session reliability coefficients observed here for SCRs were comparable to those reported in previous work (Fredrikson, Annas, Georgiades, Hursti, & Tersman, 1993; see supplementary file for an overview of studies in the field in Klingelhöfer-Jens et al., 2022) and are, as observed previously, markedly higher for individual stimuli (i.e., CSs and GSs) as opposed to CS discrimination. As the latter is a difference score, this does, however, not come as a surprise (Infantolino, Luking, Sauder, Curtin, & Hajcak, 2018; Lynam, Hoyle, & Newman, 2006; Moriarity & Alloy, 2021).

The Cronbach’s alpha coefficient for the complete CTQ-SF questionnaire was 0.89, which is consistent with previous studies (alpha = 0.94, Wingenfeld et al., 2010) and 0.85 and 0.83, for composite scales abuse and neglect, respectively.

Supplementary Table 1: Split-half reliability for SCRs

| **Phase** | **Stimulus type** | **r** | **r lower CI** | **r upper CI** |
| --- | --- | --- | --- | --- |
| **Acquisition** | CS+ | 0.802 | 0.783 | 0.820 |
|  | CS- | 0.735 | 0.710 | 0.758 |
|  | CS discr. | 0.322 | 0.274 | 0.368 |
| **Generalization** | CS+ | 0.720 | 0.693 | 0.744 |
|  | CS- | 0.470 | 0.429 | 0.510 |
|  | CS discr. | 0.346 | 0.299 | 0.391 |
|  | GS1 | 0.656 | 0.625 | 0.684 |
|  | GS2 | 0.550 | 0.512 | 0.585 |
|  | GS3 | 0.561 | 0.524 | 0.596 |
|  | GS4 | 0.288 | 0.240 | 0.336 |
| Note. CS = conditioned stimulus, GS = generalization stimulus, CS discr. = CS discrimination. | | | | |

#### Supplementary statistical analyses (main effects of task).

To test for successful manipulation during acquisition training, two-tailed paired t-tests were performed for SCRs and ratings comparing CS+ and CS- responses averaged across trials excluding the first acquisition trial as no learning could possibly have taken place due to the delay conditioning paradigm (Lonsdorf et al., 2017). To test for successful manipulation during the generalization phase, a one-way ANOVA with stimulus type as within-subject factor was calculated for SCRs and ratings. Post hoc two-tailed paired t-tests were conducted to test for a gradual increase in responses with increasing similarity to the CS+.

#### Supplementary Results: Main Effects of Task.

As published previously (Herzog et al., 2021; Schiele, Reinhard, et al., 2016), successful fear acquisition could be confirmed by significantly stronger responses to the CS+ as compared to CS- in all outcome measures (SCRs: *t*(1401) = 13.86, *p* < .001, *d* = 0.37; arousal ratings: *t*(1401) = 48.26, *p* < .001, *d* = 1.29; valence ratings: *t*(1401) = 35.57, *p* < .001, *d* = 0.95; contingency ratings: *t*(1401) = 61.00, *p* < .001, *d* = 1.63). Similarly, ANOVAs indicated significant differences between responses to the CS+, CS- and GSs during generalization phase (SCRs: *F*(4.49, 6284.85) = 60.66, *p* < .001, $\eta_{p}^{2}$ = .04; arousal ratings: *F*(3.27, 4578.72) = 1031.46, *p* < .001, $\eta_{p}^{2}$ = .42; valence ratings: *F*(3.43, 4809.75) = 766.44, *p* < .001, $\eta_{p}^{2}$ = .35; contingency ratings: *F*(3.20, 4482.78) = 2222.76, *p* < .001, $\eta_{p}^{2}$ = .61). Post-hoc tests yielded that all CSs and GSs differed from each other during generalization phase in all outcome measures (all *p*’s < 0.01), except for the comparisons of CS- vs. GS4 and GS1 vs. GS2 which were not significant (both *p*’s = 0.12).

### Supplementary robustness analyses

Robustness analyses were performed for main SCR analyses of CS discrimination during both the acquisition training and generalization phase, as well as the general reactivity by repeating all main analyses as described in the main manuscript with a) exclusion of physiological non-responders (i.e., participants with only SCRs = 0), b) exclusion of extreme outliers (i.e., data points +/- 3 x interquartile range (IQR) above/below Q3/Q1), c) study site as a covariate and d) square root transformed instead of log-transformed and range-corrected SCRs.

The robustness analyses yielded that results did not change substantially (i.e., all statistically significant results remained significant [data not shown] - with two exceptions: First, entering site as a covariate in analyses of CS discrimination during acquisition training attenuated the p-value of the childhood adversity exposure effect from *p* = 0.02 to *p* = 0.09. Likewise, the difference in CS discrimination during acquisition training between exposed and unexposed individuals in SCRs dropped from *p* = 0.02 to *p* = 0.06 when SCRs were square-root transformed instead of log-transformed and range-corrected.

### Additional figures and tables referred to in the main manuscript

Supplementary Table 2. Results of the repetition of our main analyses using linear mixed models for SCR (A), arousal (B), valence (C), and contingency ratings (D)

A)

|  | **CS_discrimination_SCR_during_ACQ** | | | | **CS_discrimination_SCR_during_GEN** | | | | **LDS_SCR** | | | | **General_reactivity_SCR** | | | |
| --- | --- | --- | --- | --- | --- | --- | --- | --- | --- | --- | --- | --- | --- | --- | --- | --- |
| *Predictors* | *Estimates* | *CI* | *p* | *df* | *Estimates* | *CI* | *p* | *df* | *Estimates* | *CI* | *p* | *df* | *Estimates* | *CI* | *p* | *df* |
| (Intercept) | 0.18 | -0.03 – 0.40 | 0.100 | 1367.26 | 0.03 | -0.25 – 0.30 | 0.841 | 1347.55 | 0.02 | -0.12 – 0.16 | 0.771 | 1387.92 | 0.16 | -0.10 – 0.43 | 0.230 | 130.64 |
| Age | 0.00 | -0.00 – 0.00 | 0.327 | 1388.92 | -0.00 | -0.00 – 0.00 | 0.334 | 1388.97 | -0.00 | -0.00 – 0.00 | 0.655 | 1163.35 | -0.00 | -0.00 – 0.00 | 0.429 | 1386.32 |
| Sex [1] | -0.02 | -0.03 – -0.01 | **<0.001** | 1388.87 | -0.00 | -0.02 – 0.01 | 0.818 | 1388.99 | 0.00 | -0.00 – 0.01 | 0.427 | 1074.14 | 0.01 | -0.00 – 0.02 | 0.176 | 1386.33 |
| dummy(School level)1 | 0.12 | -0.18 – 0.41 | 0.448 | 1386.00 | 0.24 | -0.14 – 0.63 | 0.220 | 1386.00 | 0.09 | -0.11 – 0.28 | 0.382 | 1386.16 | 0.02 | -0.32 – 0.36 | 0.913 | 1386.00 |
| dummy(School level)2 | -0.15 | -0.37 – 0.07 | 0.193 | 1386.70 | 0.03 | -0.26 – 0.31 | 0.844 | 1386.54 | 0.02 | -0.12 – 0.17 | 0.753 | 1386.60 | -0.05 | -0.30 – 0.20 | 0.706 | 1386.03 |
| dummy(School level)3 | -0.09 | -0.33 – 0.14 | 0.431 | 1386.54 | -0.06 | -0.36 – 0.24 | 0.699 | 1386.42 | 0.01 | -0.14 – 0.16 | 0.931 | 1388.10 | -0.06 | -0.33 – 0.20 | 0.654 | 1386.02 |
| dummy(School level)4 | -0.16 | -0.37 – 0.06 | 0.153 | 1386.67 | 0.04 | -0.23 – 0.31 | 0.775 | 1386.52 | -0.00 | -0.14 – 0.13 | 0.951 | 1387.75 | -0.03 | -0.27 – 0.22 | 0.830 | 1386.03 |
| dummy(School level)5 | -0.17 | -0.39 – 0.05 | 0.132 | 1386.36 | 0.05 | -0.23 – 0.33 | 0.708 | 1386.28 | -0.01 | -0.16 – 0.13 | 0.846 | 1388.84 | -0.04 | -0.29 – 0.21 | 0.754 | 1386.02 |
| dummy(School level)6 | -0.13 | -0.35 – 0.08 | 0.221 | 1386.47 | 0.02 | -0.25 – 0.30 | 0.871 | 1386.36 | -0.01 | -0.14 – 0.13 | 0.936 | 1388.97 | -0.04 | -0.28 – 0.20 | 0.751 | 1386.02 |
| dummy(School level)7 | -0.14 | -0.35 – 0.07 | 0.193 | 1386.45 | 0.05 | -0.23 – 0.32 | 0.745 | 1386.34 | -0.01 | -0.15 – 0.13 | 0.898 | 1388.96 | -0.04 | -0.28 – 0.20 | 0.754 | 1386.02 |
| dummy(School level)8 | -0.14 | -0.36 – 0.07 | 0.181 | 1386.38 | 0.03 | -0.24 – 0.31 | 0.809 | 1386.29 | -0.01 | -0.14 – 0.13 | 0.928 | 1388.97 | -0.05 | -0.29 – 0.20 | 0.713 | 1386.02 |
| dummy(School level)9 | -0.21 | -0.51 – 0.09 | 0.163 | 1386.59 | -0.09 | -0.47 – 0.30 | 0.659 | 1386.45 | -0.10 | -0.30 – 0.09 | 0.290 | 1387.60 | -0.05 | -0.40 – 0.29 | 0.754 | 1386.03 |
| Childhood adversity [1] | -0.02 | -0.04 – -0.00 | **0.017** | 1386.71 | -0.02 | -0.04 – -0.00 | **0.024** | 1386.55 | -0.01 | -0.02 – 0.00 | 0.178 | 1386.58 | -0.02 | -0.04 – -0.00 | **0.024** | 1386.03 |
| **Random Effects** | | | | | | | | | | | | | | | | |
| σ^2^ | 0.01 | | | | 0.02 | | | | 0.00 | | | | 0.02 | | | |
| τ_00_ | 0.00 _Site_ | | | | 0.00 _Site_ | | | | 0.00 _Site_ | | | | 0.01 _Site_ | | | |
| ICC | 0.03 | | | | 0.04 | | | |  | | | | 0.41 | | | |
| N | 4 _Site_ | | | | 4 _Site_ | | | | 4 _Site_ | | | | 4 _Site_ | | | |
| Observations | 1402 | | | | 1402 | | | | 1402 | | | | 1402 | | | |
| Marginal R^2^ / Conditional R^2^ | 0.021 / 0.049 | | | | 0.012 / 0.049 | | | | 0.006 / NA | | | | 0.004 / 0.414 | | | |

B)

|  | **CS_discrimination_arousal_ratings_during_ACQ** | | | | **CS_discrimination_arousal_ratings_during_GEN** | | | | **LDS_arousal_ratings** | | | | **General_reactivity_arousal_ratings** | | | |
| --- | --- | --- | --- | --- | --- | --- | --- | --- | --- | --- | --- | --- | --- | --- | --- | --- |
| *Predictors* | *Estimates* | *CI* | *p* | *df* | *Estimates* | *CI* | *p* | *df* | *Estimates* | *CI* | *p* | *df* | *Estimates* | *CI* | *p* | *df* |
| (Intercept) | 4.73 | 0.46 – 9.00 | **0.030** | 1387.12 | 3.89 | -0.92 – 8.69 | 0.113 | 1386.01 | 1.37 | -1.27 – 4.01 | 0.308 | 1388.45 | 3.22 | 0.85 – 5.59 | **0.008** | 1333.36 |
| Age | -0.02 | -0.04 – 0.01 | 0.190 | 1381.61 | 0.00 | -0.02 – 0.03 | 0.854 | 1384.27 | 0.01 | -0.00 – 0.03 | 0.063 | 1213.66 | -0.01 | -0.03 – -0.00 | **0.034** | 1388.87 |
| Sex [1] | -0.72 | -0.96 – -0.49 | **<0.001** | 1380.21 | -0.68 | -0.94 – -0.42 | **<0.001** | 1383.34 | -0.10 | -0.24 – 0.05 | 0.198 | 1151.26 | -0.22 | -0.35 – -0.09 | **0.001** | 1388.90 |
| dummy(School level)1 | -2.02 | -7.98 – 3.95 | 0.507 | 1386.01 | 1.00 | -5.70 – 7.70 | 0.769 | 1386.01 | -1.49 | -5.17 – 2.19 | 0.428 | 1386.12 | 1.74 | -1.56 – 5.03 | 0.301 | 1386.00 |
| dummy(School level)2 | -1.93 | -6.33 – 2.48 | 0.391 | 1387.53 | -1.08 | -6.03 – 3.87 | 0.668 | 1387.35 | -1.20 | -3.92 – 1.52 | 0.388 | 1388.08 | 0.66 | -1.78 – 3.09 | 0.596 | 1386.48 |
| dummy(School level)3 | -1.96 | -6.58 – 2.67 | 0.407 | 1387.23 | -1.29 | -6.49 – 3.91 | 0.627 | 1387.07 | -1.02 | -3.88 – 1.84 | 0.484 | 1388.80 | 0.45 | -2.11 – 3.01 | 0.730 | 1386.37 |
| dummy(School level)4 | -1.46 | -5.71 – 2.79 | 0.499 | 1387.45 | -0.86 | -5.64 – 3.91 | 0.723 | 1387.28 | -1.20 | -3.82 – 1.43 | 0.372 | 1388.60 | 1.26 | -1.09 – 3.61 | 0.294 | 1386.46 |
| dummy(School level)5 | -0.71 | -5.06 – 3.64 | 0.747 | 1386.81 | -0.42 | -5.31 – 4.47 | 0.868 | 1386.71 | -1.80 | -4.49 – 0.89 | 0.189 | 1388.62 | 1.57 | -0.83 – 3.98 | 0.200 | 1386.25 |
| dummy(School level)6 | -0.80 | -5.06 – 3.46 | 0.712 | 1387.04 | -0.48 | -5.26 – 4.31 | 0.845 | 1386.91 | -1.34 | -3.97 – 1.29 | 0.317 | 1388.98 | 1.16 | -1.19 – 3.51 | 0.333 | 1386.32 |
| dummy(School level)7 | -1.27 | -5.49 – 2.96 | 0.556 | 1387.01 | -0.47 | -5.22 – 4.27 | 0.845 | 1386.89 | -1.19 | -3.80 – 1.42 | 0.373 | 1388.98 | 1.28 | -1.05 – 3.62 | 0.281 | 1386.30 |
| dummy(School level)8 | -1.29 | -5.52 – 2.93 | 0.549 | 1386.85 | -0.42 | -5.17 – 4.33 | 0.862 | 1386.75 | -1.23 | -3.84 – 1.38 | 0.356 | 1388.80 | 1.29 | -1.04 – 3.63 | 0.278 | 1386.26 |
| dummy(School level)9 | 1.99 | -3.99 – 7.96 | 0.514 | 1387.32 | 2.92 | -3.79 – 9.64 | 0.393 | 1387.16 | -3.15 | -6.84 – 0.55 | 0.095 | 1388.59 | 1.59 | -1.72 – 4.89 | 0.346 | 1386.40 |
| Childhood adversity [1] | 0.26 | -0.06 – 0.59 | 0.108 | 1387.55 | 0.06 | -0.30 – 0.42 | 0.752 | 1387.37 | 0.04 | -0.16 – 0.24 | 0.670 | 1388.05 | 0.03 | -0.15 – 0.21 | 0.738 | 1386.48 |
| **Random Effects** | | | | | | | | | | | | | | | | |
| σ^2^ | 4.62 | | | | 5.83 | | | | 1.76 | | | | 1.41 | | | |
| τ_00_ | 0.05 _Site_ | | | | 0.08 _Site_ | | | | 0.00 _Site_ | | | | 0.06 _Site_ | | | |
| ICC | 0.01 | | | | 0.01 | | | | 0.00 | | | | 0.04 | | | |
| N | 4 _Site_ | | | | 4 _Site_ | | | | 4 _Site_ | | | | 4 _Site_ | | | |
| Observations | 1402 | | | | 1402 | | | | 1402 | | | | 1402 | | | |
| Marginal R^2^ / Conditional R^2^ | 0.036 / 0.046 | | | | 0.021 / 0.034 | | | | 0.009 / 0.010 | | | | 0.019 / 0.061 | | | |

C)

|  | **CS_discrimination_valence_ratings_during_ACQ** | | | | **CS_discrimination_valence_ratings_during_GEN** | | | | **LDS_valence_ratings** | | | | **General_reactivity_valence_ratings** | | | |
| --- | --- | --- | --- | --- | --- | --- | --- | --- | --- | --- | --- | --- | --- | --- | --- | --- |
| *Predictors* | *Estimates* | *CI* | *p* | *df* | *Estimates* | *CI* | *p* | *df* | *Estimates* | *CI* | *p* | *df* | *Estimates* | *CI* | *p* | *df* |
| (Intercept) | 3.58 | -0.74 – 7.90 | 0.104 | 1377.81 | 3.40 | -1.67 – 8.48 | 0.189 | 1388.66 | 2.39 | -0.25 – 5.02 | 0.076 | 1382.58 | 5.82 | 3.94 – 7.70 | **<0.001** | 1365.32 |
| Age | -0.03 | -0.06 – -0.01 | **0.005** | 1388.31 | -0.02 | -0.04 – 0.01 | 0.266 | 1243.67 | 0.00 | -0.01 – 0.02 | 0.884 | 1387.16 | -0.01 | -0.02 – 0.00 | 0.062 | 1388.96 |
| Sex [1] | -0.55 | -0.79 – -0.31 | **<0.001** | 1388.11 | -0.66 | -0.94 – -0.38 | **<0.001** | 1195.80 | -0.07 | -0.21 – 0.08 | 0.352 | 1386.74 | -0.01 | -0.11 – 0.10 | 0.916 | 1388.92 |
| dummy(School level)1 | -2.03 | -8.05 – 3.98 | 0.507 | 1386.00 | -1.02 | -8.11 – 6.08 | 0.779 | 1386.10 | -1.75 | -5.42 – 1.93 | 0.351 | 1386.00 | -0.84 | -3.46 – 1.77 | 0.527 | 1386.00 |
| dummy(School level)2 | -2.53 | -6.98 – 1.91 | 0.263 | 1386.89 | -1.19 | -6.44 – 4.05 | 0.655 | 1388.63 | -2.08 | -4.79 – 0.64 | 0.134 | 1387.08 | -1.14 | -3.07 – 0.79 | 0.246 | 1386.68 |
| dummy(School level)3 | -0.48 | -5.15 – 4.19 | 0.840 | 1386.70 | -1.20 | -6.71 – 4.30 | 0.668 | 1388.97 | -1.33 | -4.18 – 1.52 | 0.361 | 1386.84 | -1.55 | -3.58 – 0.48 | 0.134 | 1386.52 |
| dummy(School level)4 | -0.31 | -4.59 – 3.98 | 0.889 | 1386.85 | 0.19 | -4.87 – 5.25 | 0.941 | 1388.88 | -1.91 | -4.53 – 0.71 | 0.154 | 1387.02 | -0.85 | -2.72 – 1.01 | 0.371 | 1386.65 |
| dummy(School level)5 | 0.07 | -4.32 – 4.46 | 0.976 | 1386.46 | 0.58 | -4.60 – 5.75 | 0.827 | 1388.45 | -2.30 | -4.99 – 0.38 | 0.092 | 1386.56 | -0.36 | -2.27 – 1.55 | 0.711 | 1386.35 |
| dummy(School level)6 | 0.07 | -4.22 – 4.37 | 0.973 | 1386.60 | 0.11 | -4.95 – 5.18 | 0.965 | 1388.89 | -1.81 | -4.44 – 0.81 | 0.176 | 1386.72 | -0.91 | -2.78 – 0.96 | 0.338 | 1386.45 |
| dummy(School level)7 | -0.53 | -4.79 – 3.73 | 0.807 | 1386.57 | -0.12 | -5.14 – 4.91 | 0.964 | 1388.89 | -1.85 | -4.46 – 0.75 | 0.163 | 1386.70 | -0.74 | -2.59 – 1.11 | 0.433 | 1386.43 |
| dummy(School level)8 | -0.40 | -4.67 – 3.86 | 0.853 | 1386.48 | 0.04 | -4.99 – 5.07 | 0.987 | 1388.64 | -1.83 | -4.43 – 0.78 | 0.170 | 1386.59 | -0.75 | -2.61 – 1.10 | 0.426 | 1386.36 |
| dummy(School level)9 | 2.20 | -3.83 – 8.23 | 0.474 | 1386.75 | 2.68 | -4.43 – 9.80 | 0.459 | 1388.90 | -3.61 | -7.30 – 0.07 | 0.055 | 1386.91 | -0.41 | -3.04 – 2.21 | 0.757 | 1386.57 |
| Childhood adversity [1] | -0.02 | -0.35 – 0.31 | 0.902 | 1386.90 | -0.01 | -0.40 – 0.37 | 0.950 | 1388.60 | -0.03 | -0.23 – 0.17 | 0.754 | 1387.09 | -0.04 | -0.18 – 0.10 | 0.581 | 1386.69 |
| **Random Effects** | | | | | | | | | | | | | | | | |
| σ^2^ | 4.70 | | | | 6.53 | | | | 1.76 | | | | 0.89 | | | |
| τ_00_ | 0.10 _Site_ | | | | 0.01 _Site_ | | | | 0.03 _Site_ | | | | 0.03 _Site_ | | | |
| ICC | 0.02 | | | | 0.00 | | | | 0.02 | | | | 0.03 | | | |
| N | 4 _Site_ | | | | 4 _Site_ | | | | 4 _Site_ | | | | 4 _Site_ | | | |
| Observations | 1402 | | | | 1402 | | | | 1402 | | | | 1402 | | | |
| Marginal R^2^ / Conditional R^2^ | 0.035 / 0.055 | | | | 0.021 / 0.022 | | | | 0.006 / 0.023 | | | | 0.015 / 0.044 | | | |

D)

|  | **CS_discrimination_contingency_ratings_during_ACQ** | | | | **CS_discrimination_contingency_ratings_during_GEN** | | | | **LDS_contingency_ratings** | | | | **General_reactivity_contingency_ratings** | | | |
| --- | --- | --- | --- | --- | --- | --- | --- | --- | --- | --- | --- | --- | --- | --- | --- | --- |
| *Predictors* | *Estimates* | *CI* | *p* | *df* | *Estimates* | *CI* | *p* | *df* | *Estimates* | *CI* | *p* | *df* | *Estimates* | *CI* | *p* | *df* |
| (Intercept) | 5.56 | -57.39 – 68.52 | 0.862 | 1381.99 | 83.30 | 25.26 – 141.34 | **0.005** | 1384.67 | 43.25 | 10.35 – 76.14 | **0.010** | 1379.65 | 21.72 | -3.20 – 46.65 | 0.088 | 1169.04 |
| Age | -0.18 | -0.53 – 0.16 | 0.303 | 1387.39 | 0.20 | -0.12 – 0.52 | 0.217 | 1385.88 | 0.03 | -0.15 – 0.21 | 0.728 | 1388.01 | 0.00 | -0.15 – 0.15 | 0.963 | 1170.37 |
| Sex [1] | -6.28 | -9.74 – -2.82 | **<0.001** | 1387.01 | -3.72 | -6.91 – -0.53 | **0.022** | 1385.23 | 0.58 | -1.22 – 2.39 | 0.526 | 1387.75 | -1.58 | -3.05 – -0.11 | **0.036** | 1159.35 |
| dummy(School level)1 | 94.82 | 7.05 – 182.59 | **0.034** | 1386.00 | 0.20 | -80.74 – 81.14 | 0.996 | 1386.01 | 0.03 | -45.82 – 45.88 | 0.999 | 1386.00 | 9.17 | -25.54 – 43.88 | 0.604 | 1169.00 |
| dummy(School level)2 | 21.75 | -43.07 – 86.58 | 0.510 | 1387.05 | -45.11 | -104.90 – 14.67 | 0.139 | 1387.21 | -30.03 | -63.89 – 3.83 | 0.082 | 1386.95 | 9.97 | -15.78 – 35.73 | 0.447 | 1170.68 |
| dummy(School level)3 | 35.78 | -32.32 – 103.88 | 0.303 | 1386.82 | -34.55 | -97.35 – 28.25 | 0.281 | 1386.96 | -29.19 | -64.76 – 6.38 | 0.108 | 1386.74 | 13.87 | -13.07 – 40.81 | 0.313 | 1170.47 |
| dummy(School level)4 | 46.02 | -16.56 – 108.60 | 0.149 | 1386.99 | -32.68 | -90.40 – 25.03 | 0.267 | 1387.15 | -29.54 | -62.23 – 3.15 | 0.076 | 1386.90 | 20.52 | -4.28 – 45.31 | 0.105 | 1170.59 |
| dummy(School level)5 | 55.51 | -8.54 – 119.57 | 0.089 | 1386.54 | -35.90 | -94.97 – 23.17 | 0.233 | 1386.63 | -35.79 | -69.25 – -2.33 | **0.036** | 1386.49 | 23.91 | -1.66 – 49.49 | 0.067 | 1169.58 |
| dummy(School level)6 | 49.68 | -12.99 – 112.35 | 0.120 | 1386.70 | -31.66 | -89.46 – 26.13 | 0.283 | 1386.82 | -32.52 | -65.26 – 0.22 | 0.052 | 1386.64 | 21.10 | -3.75 – 45.94 | 0.096 | 1170.10 |
| dummy(School level)7 | 53.68 | -8.52 – 115.88 | 0.091 | 1386.68 | -29.33 | -86.69 – 28.03 | 0.316 | 1386.79 | -30.67 | -63.16 – 1.82 | 0.064 | 1386.61 | 18.22 | -6.39 – 42.83 | 0.147 | 1170.18 |
| dummy(School level)8 | 56.22 | -6.02 – 118.46 | 0.077 | 1386.57 | -30.69 | -88.08 – 26.71 | 0.294 | 1386.67 | -30.51 | -63.02 – 2.00 | 0.066 | 1386.51 | 17.44 | -7.18 – 42.07 | 0.165 | 1169.91 |
| dummy(School level)9 | 89.88 | 1.91 – 177.85 | **0.045** | 1386.89 | -57.56 | -138.69 – 23.57 | 0.164 | 1387.04 | -35.92 | -81.87 – 10.03 | 0.125 | 1386.80 | -0.23 | -35.05 – 34.58 | 0.989 | 1170.57 |
| Childhood adversity [1] | -0.86 | -5.61 – 3.89 | 0.723 | 1387.06 | -1.26 | -5.65 – 3.12 | 0.572 | 1387.23 | 1.13 | -1.35 – 3.62 | 0.370 | 1386.96 | 0.77 | -1.27 – 2.81 | 0.461 | 1170.80 |
| **Random Effects** | | | | | | | | | | | | | | | | |
| σ^2^ | 1001.00 | | | | 851.30 | | | | 273.12 | | | | 156.50 | | | |
| τ_00_ | 18.22 _Site_ | | | | 12.82 _Site_ | | | | 5.58 _Site_ | | | | 0.88 _Site_ | | | |
| ICC | 0.02 | | | | 0.01 | | | | 0.02 | | | | 0.01 | | | |
| N | 4 _Site_ | | | | 4 _Site_ | | | | 4 _Site_ | | | | 3 _Site_ | | | |
| Observations | 1402 | | | | 1402 | | | | 1402 | | | | 1184 | | | |
| Marginal R^2^ / Conditional R^2^ | 0.029 / 0.047 | | | | 0.009 / 0.024 | | | | 0.008 / 0.028 | | | | 0.020 / 0.025 | | | |

*Note.* Due to its categorical nature, we included school level as a dummy variable.

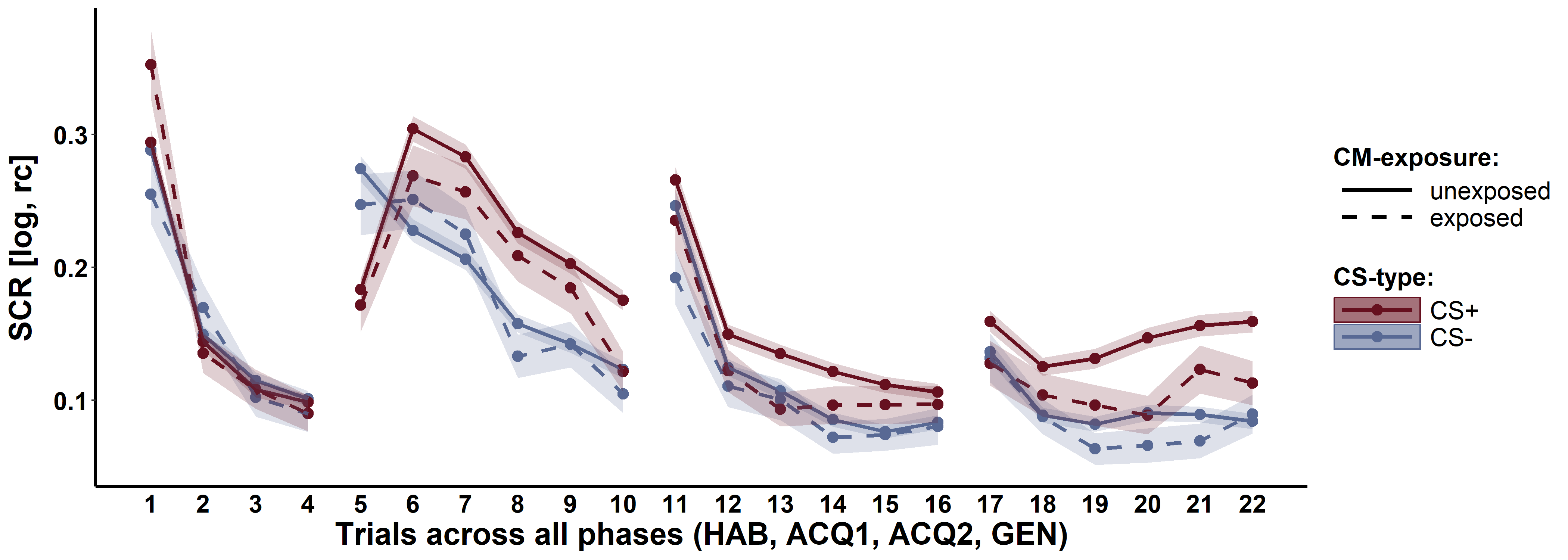

Supplementary Figure 4: Trial-by-trial SCR data across all experimental phases for the CS+ (red) and CS- (blue) for individuals exposed (dashed lines) and unexposed (solid lines) to childhood adversity separately. Ribbons represent standard errors of the means (SEMs) including n_unexposed_ = 1199 and n_exposed_ = 203.

Supplementary Table 3: Exploratory results of testing the cumulative risk model involving severity groups using ANOVAs with exposure to abuse as between-subject factor and CS discrimination, LDS, and general reactivity as dependent variable

| **Outcome** | **Phase** | **Measure** | **Mean 'none'** | **SD 'none'** | **Mean 'low'** | **SD 'low'** | **Mean 'moderate'** | **SD 'moderate'** | **Mean 'severe'** | **SD 'severe'** | ***df_Num_*** | ***df_Den_*** | ***F*** | ***p*** | ***partial Eta^2^*** |
| --- | --- | --- | --- | --- | --- | --- | --- | --- | --- | --- | --- | --- | --- | --- | --- |
| **CS discrimination** | **ACQ** | **SCR** | 0.05 | 0.11 | 0.04 | 0.10 | 0.02 | 0.11 | 0.03 | 0.10 | 3 | 1,398 | 2.35 | 0.071 | 0.00 |
|  |  | **Arousal ratings** | 2.80 | 2.21 | 2.76 | 2.18 | 2.99 | 2.07 | 3.13 | 2.21 | 3 | 1,398 | 0.86 | 0.459 | 0.00 |
|  |  | **Valence ratings** | 2.05 | 2.19 | 2.21 | 2.21 | 2.11 | 2.21 | 2.00 | 2.41 | 3 | 1,398 | 0.54 | 0.655 | 0.00 |
|  |  | **Contingency ratings** | 53.01 | 31.88 | 52.06 | 32.00 | 52.37 | 33.16 | 48.03 | 35.02 | 3 | 1,398 | 0.51 | 0.673 | 0.00 |
|  | **GEN** | **SCR** | 0.05 | 0.14 | 0.05 | 0.14 | 0.01 | 0.12 | 0.04 | 0.14 | 3 | 1,398 | 2.36 | 0.070 | 0.00 |
|  |  | **Arousal ratings** | 3.23 | 2.44 | 3.02 | 2.42 | 3.12 | 2.35 | 3.39 | 2.77 | 3 | 1,398 | 0.91 | 0.437 | 0.00 |
|  |  | **Valence ratings** | 2.66 | 2.64 | 2.73 | 2.41 | 2.66 | 2.53 | 2.71 | 2.89 | 3 | 1,398 | 0.07 | 0.975 | 0.00 |
|  |  | **Contingency ratings** | *58.20* | *28.95* | *56.12* | *28.66* | *56.93* | *30.43* | *54.55* | *35.09* | *3* | *137* | *0.80* | *0.498* | *0.13* |
| **LDS** | **GEN** | **SCR** | 0.01 | 0.07 | 0.01 | 0.07 | 0.00 | 0.06 | 0.00 | 0.06 | 3 | 1,398 | 0.60 | 0.615 | 0.00 |
|  |  | **Arousal ratings** | 0.52 | 1.32 | 0.38 | 1.32 | 0.48 | 1.40 | 0.63 | 1.25 | 3 | 1,398 | 1.38 | 0.248 | 0.00 |
|  |  | **Valence ratings** | 0.56 | 1.33 | 0.54 | 1.33 | 0.50 | 1.23 | 0.59 | 1.49 | 3 | 1,398 | 0.14 | 0.935 | 0.00 |
|  |  | **Contingency ratings** | 14.11 | 16.42 | 13.91 | 16.79 | 14.82 | 17.43 | 16.10 | 16.32 | 3 | 1,398 | 0.40 | 0.754 | 0.00 |
| **General reactivity** | **ALL** | **SCR** | 0.10 | 0.16 | 0.09 | 0.12 | 0.07 | 0.10 | 0.07 | 0.10 | 3 | 1,398 | 1.64 | 0.178 | 0.00 |
|  |  | **Arousal ratings** | 4.01 | 1.22 | 4.04 | 1.24 | 4.15 | 1.09 | 3.79 | 1.12 | 3 | 1,398 | 1.43 | 0.234 | 0.00 |
|  |  | **Valence ratings** | 4.81 | 0.96 | 4.77 | 0.98 | 4.78 | 0.83 | 4.64 | 0.92 | 3 | 1,398 | 0.78 | 0.508 | 0.00 |
|  |  | **Contingency ratings** | *39.42* | *12.21* | *39.09* | *13.71* | *40.54* | *11.64* | *39.50* | *11.32* | *3* | *120* | *0.54* | *0.656* | *0.15* |
| Note. ACQ = acquisition training, GEN = generalization phase, LDS = linear deviation score, SD = standard deviation. Italic lines indicate the application of robust ANOVAs. In this context, effect sizes do not indicate partial eta squared, but the explanatory measure of effect size (Mair & Wilcox, 2020). Values of 0.10, 0.30, and 0.50 represent small, medium, and large effect sizes, respectively. | | | | | | | | | | | | | | | |

Supplementary Table 4: Exploratory results of testing the cumulative risk model involving the number of subscales exceeding an at least moderate cut-off using regressions with the number of subscales as predictor and CS discrimination, LDS, and general reactivity as criterion

| **Outcome** | **Phase** | **Measure** | ***beta*** | ***SE_b_*** | ***LL (95% CI)*** | ***UL (95% CI)*** | ***Beta*** | ***t*** | ***df*** | ***p*** | ***R^2^*** | ***Cohen's f^2^*** |
| --- | --- | --- | --- | --- | --- | --- | --- | --- | --- | --- | --- | --- |
| **CS discrimination** | **ACQ** | **SCR** | **-0.01** | **0.00** | **-0.02** | **0.00** | **-0.07** | **-2.62** | **1,400** | **0.009** | **0** | **0** |
|  |  | **Arousal ratings** | 0.07 | 0.09 | -0.09 | 0.24 | 0.02 | 0.85 | 1,400 | 0.393 | 0 | 0 |
|  |  | **Valence ratings** | -0.05 | 0.09 | -0.22 | 0.12 | -0.01 | -0.55 | 1,400 | 0.582 | 0 | 0 |
|  |  | **Contingency ratings** | **-2.53** | **1.25** | **-4.98** | **-0.07** | **-0.05** | **-2.02** | **1,400** | **0.044** | **0** | **0** |
|  | **GEN** | **SCR** | -0.01 | 0.00 | -0.02 | 0.00 | -0.03 | -1.23 | 1,400 | 0.218 | 0 | 0 |
|  |  | **Arousal ratings** | 0.00 | 0.10 | -0.19 | 0.19 | 0.00 | 0.00 | 1,400 | 0.997 | 0 | 0 |
|  |  | **Valence ratings** | 0.03 | 0.10 | -0.17 | 0.22 | 0.01 | 0.27 | 1,400 | 0.789 | 0 | 0 |
|  |  | **Contingency ratings** | -1.61 | 1.14 | -3.85 | 0.63 | -0.04 | -1.41 | 1,400 | 0.159 | 0 | 0 |
| **LDS** | **GEN** | **SCR** | 0.00 | 0.00 | -0.01 | 0.00 | -0.01 | -0.48 | 1,400 | 0.629 | 0 | 0 |
|  |  | **Arousal ratings** | 0.03 | 0.05 | -0.07 | 0.13 | 0.01 | 0.56 | 1,400 | 0.579 | 0 | 0 |
|  |  | **Valence ratings** | 0.00 | 0.05 | -0.10 | 0.10 | 0.00 | 0.04 | 1,400 | 0.965 | 0 | 0 |
|  |  | **Contingency ratings** | 0.71 | 0.65 | -0.56 | 1.98 | 0.03 | 1.10 | 1,400 | 0.272 | 0 | 0 |
| **General reactivity** | **ALL** | **SCR** | -0.01 | 0.00 | -0.02 | 0.00 | -0.04 | -1.91 | 1,400 | 0.057 | 0 | 0 |
|  |  | **Arousal ratings** | -0.03 | 0.05 | -0.12 | 0.06 | -0.02 | -0.65 | 1,400 | 0.517 | 0 | 0 |
|  |  | **Valence ratings** | -0.04 | 0.04 | -0.11 | 0.03 | -0.03 | -1.06 | 1,400 | 0.290 | 0 | 0 |
|  |  | **Contingency ratings** | 0.59 | 0.55 | -0.48 | 1.66 | 0.03 | 1.08 | 1,182 | 0.281 | 0 | 0 |
| Note. ACQ = acquisition training, GEN = generalization phase, LDS = linear deviation score. Bold numbers indicate significant results (p < 0.05). | | | | | | | | | | | | |

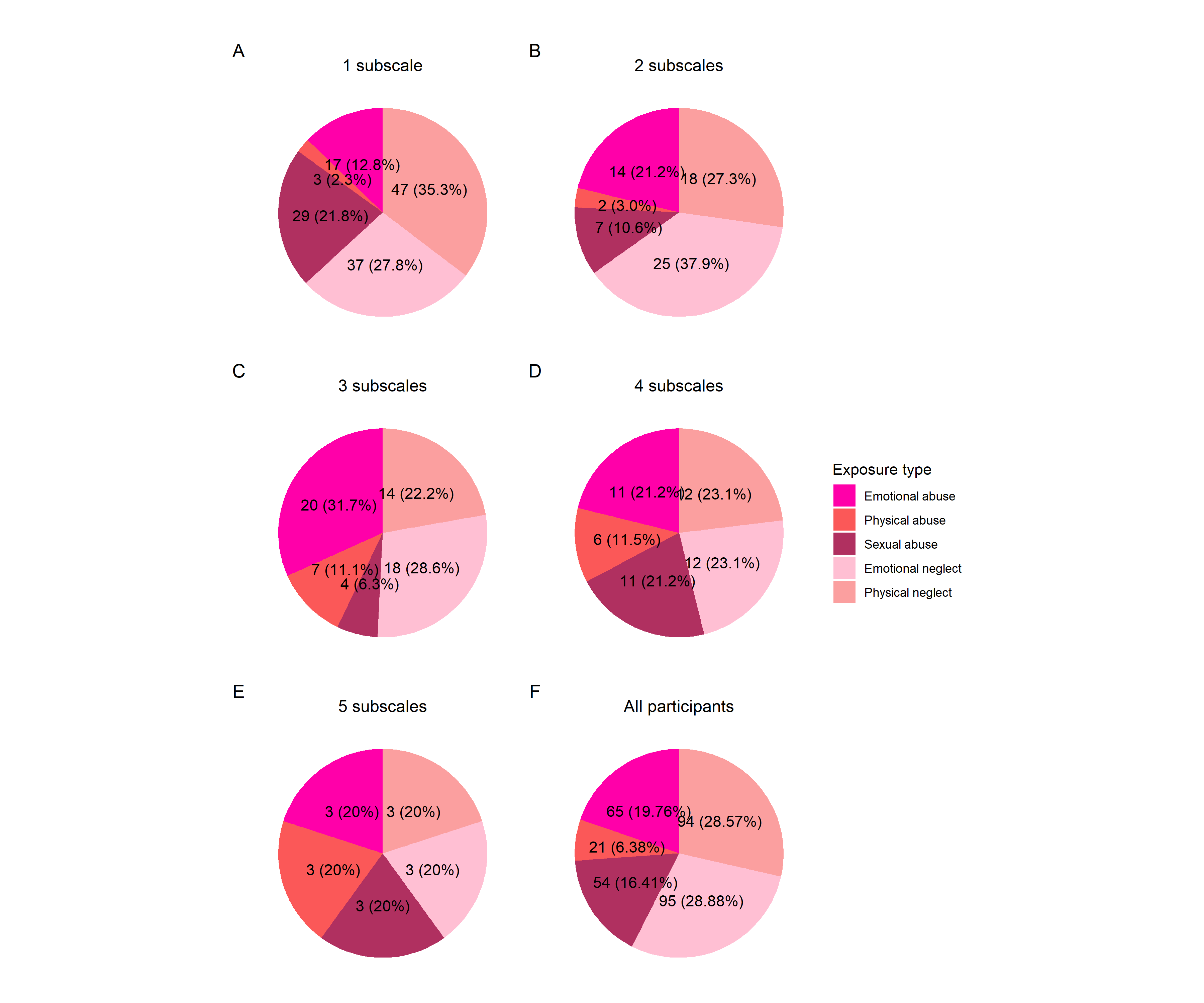

Supplementary Figure 5: Illustration of the distribution of the different numbers (i.e., 1 - 5) of exceeded subscales among the CTQ exposure types emotional abuse, physical abuse, emotional neglect, and physical neglect (A - E) and distribution of exposure types across all participants (F).

Supplementary Table 5: Exploratory results of testing the specificity model using regressions with exposure to abuse as predictor and CS discrimination, LDS, and general reactivity as criterion

| **Outcome** | **Phase** | **Measure** | ***beta*** | ***SE_b_*** | ***LL (95% CI)*** | ***UL (95% CI)*** | ***Beta*** | ***t*** | ***df*** | ***p*** | ***R^2^*** | ***Cohen's f^2^*** |
| --- | --- | --- | --- | --- | --- | --- | --- | --- | --- | --- | --- | --- |
| **CS discrimination** | **ACQ** | **SCR** | 0.00 | 0.00 | 0.00 | 0.00 | -0.03 | -1.26 | 1,400 | 0.209 | 0 | 0 |
|  |  | **Arousal ratings** | 0.01 | 0.01 | -0.02 | 0.03 | 0.02 | 0.59 | 1,400 | 0.556 | 0 | 0 |
|  |  | **Valence ratings** | 0.00 | 0.01 | -0.02 | 0.03 | 0.01 | 0.27 | 1,400 | 0.789 | 0 | 0 |
|  |  | **Contingency ratings** | -0.25 | 0.20 | -0.64 | 0.14 | -0.03 | -1.25 | 1,400 | 0.211 | 0 | 0 |
|  | **GEN** | **SCR** | 0.00 | 0.00 | 0.00 | 0.00 | 0.03 | 1.01 | 1,400 | 0.311 | 0 | 0 |
|  |  | **Arousal ratings** | 0.00 | 0.01 | -0.02 | 0.04 | 0.01 | 0.35 | 1,400 | 0.725 | 0 | 0 |
|  |  | **Valence ratings** | 0.02 | 0.02 | -0.01 | 0.05 | 0.03 | 1.20 | 1,400 | 0.230 | 0 | 0 |
|  |  | **Contingency ratings** | -0.13 | 0.18 | -0.48 | 0.23 | -0.02 | -0.71 | 1,400 | 0.479 | 0 | 0 |
| **LDS** | **GEN** | **SCR** | 0.00 | 0.00 | 0.00 | 0.00 | -0.01 | -0.22 | 1,400 | 0.825 | 0 | 0 |
|  |  | **Arousal ratings** | 0.00 | 0.01 | -0.02 | 0.01 | 0.00 | -0.08 | 1,400 | 0.933 | 0 | 0 |
|  |  | **Valence ratings** | 0.00 | 0.01 | -0.02 | 0.01 | -0.01 | -0.47 | 1,400 | 0.639 | 0 | 0 |
|  |  | **Contingency ratings** | 0.07 | 0.10 | -0.13 | 0.27 | 0.02 | 0.68 | 1,400 | 0.496 | 0 | 0 |
| **General reactivity** | **ALL** | **SCR** | 0.00 | 0.00 | 0.00 | 0.00 | 0.00 | -0.02 | 1,400 | 0.984 | 0 | 0 |
|  |  | **Arousal ratings** | 0.00 | 0.01 | -0.01 | 0.02 | 0.02 | 0.66 | 1,400 | 0.511 | 0 | 0 |
|  |  | **Valence ratings** | 0.00 | 0.01 | -0.01 | 0.01 | 0.00 | -0.19 | 1,400 | 0.853 | 0 | 0 |
|  |  | **Contingency ratings** | 0.12 | 0.09 | -0.05 | 0.29 | 0.04 | 1.38 | 1,182 | 0.167 | 0 | 0 |
| Note. ACQ = acquisition training, GEN = generalization phase, LDS = linear deviation score. | | | | | | | | | | | | |

Supplementary Table 6: Exploratory results of testing the specificity model using regressions with exposure to neglect as predictor and CS discrimination, LDS, and general reactivity as criterion

| **Outcome** | **Phase** | **Measure** | ***beta*** | ***SE_b_*** | ***LL (95% CI)*** | ***UL (95% CI)*** | ***Beta*** | ***t*** | ***df*** | ***p*** | ***R^2^*** | ***Cohen's f^2^*** |
| --- | --- | --- | --- | --- | --- | --- | --- | --- | --- | --- | --- | --- |
| **CS discrimination** | **ACQ** | **SCR** | **0.00** | **0.00** | **0.00** | **0.00** | **-0.07** | **-2.53** | **1,400** | **0.012** | **0** | **0** |
|  |  | **Arousal ratings** | 0.01 | 0.01 | -0.02 | 0.03 | 0.01 | 0.50 | 1,400 | 0.615 | 0 | 0 |
|  |  | **Valence ratings** | 0.00 | 0.01 | -0.02 | 0.03 | 0.00 | 0.10 | 1,400 | 0.919 | 0 | 0 |
|  |  | **Contingency ratings** | **-0.41** | **0.17** | **-0.75** | **-0.07** | **-0.06** | **-2.36** | **1,400** | **0.018** | **0** | **0** |
|  | **GEN** | **SCR** | 0.00 | 0.00 | 0.00 | 0.00 | -0.04 | -1.29 | 1,400 | 0.196 | 0 | 0 |
|  |  | **Arousal ratings** | -0.01 | 0.01 | -0.03 | 0.02 | -0.02 | -0.59 | 1,400 | 0.558 | 0 | 0 |
|  |  | **Valence ratings** | 0.00 | 0.01 | -0.03 | 0.02 | -0.01 | -0.27 | 1,400 | 0.789 | 0 | 0 |
|  |  | **Contingency ratings** | **-0.40** | **0.16** | **-0.71** | **-0.09** | **-0.07** | **-2.52** | **1,400** | **0.012** | **0** | **0** |
| **LDS** | **GEN** | **SCR** | 0.00 | 0.00 | 0.00 | 0.00 | -0.02 | -0.66 | 1,400 | 0.512 | 0 | 0 |
|  |  | **Arousal ratings** | 0.00 | 0.01 | -0.02 | 0.01 | -0.01 | -0.25 | 1,400 | 0.799 | 0 | 0 |
|  |  | **Valence ratings** | 0.00 | 0.01 | -0.01 | 0.01 | 0.00 | 0.08 | 1,400 | 0.935 | 0 | 0 |
|  |  | **Contingency ratings** | 0.01 | 0.09 | -0.17 | 0.19 | 0.00 | 0.11 | 1,400 | 0.915 | 0 | 0 |
| **General reactivity** | **ALL** | **SCR** | **0.00** | **0.00** | **0.00** | **0.00** | **-0.06** | **-2.31** | **1,400** | **0.021** | **0** | **0** |
|  |  | **Arousal ratings** | 0.00 | 0.01 | -0.02 | 0.01 | -0.02 | -0.67 | 1,400 | 0.504 | 0 | 0 |
|  |  | **Valence ratings** | -0.01 | 0.00 | -0.02 | 0.00 | -0.05 | -1.76 | 1,400 | 0.079 | 0 | 0 |
|  |  | **Contingency ratings** | 0.07 | 0.07 | -0.08 | 0.22 | 0.03 | 0.91 | 1,182 | 0.365 | 0 | 0 |
| Note. ACQ = acquisition training, GEN = generalization phase, LDS = linear deviation score. Bold numbers indicate significant results (p < .05). | | | | | | | | | | | | |

Supplementary Table 7: Exploratory results of testing the dimensional model using multiple regressions with exposure to both abuse and neglect as predictors and CS discrimination, LDS, and general reactivity as criterion

| **Outcome** | **Phase** | **Measure** | ***predictor*** | ***beta*** | ***SE_b_*** | ***LL (95% CI)*** | ***UL (95% CI)*** | ***Beta*** | ***t*** | ***df*** | ***p*** | ***adj. R^2^*** | **Cohen's f^2^** |
| --- | --- | --- | --- | --- | --- | --- | --- | --- | --- | --- | --- | --- | --- |
| **CS discrimination** | **ACQ** | **SCR** | abuse | 0.00 | 0.00 | -0.01 | 0.00 | -0.09 | -1.19 | 1,398 | 0.234 | 0 | 0 |
|  |  |  | **neglect** | **0.00** | **0.00** | **-0.01** | **0.00** | **-0.17** | **-2.33** | **1,398** | **0.020** | **0** | 0 |
|  |  |  | interaction | 0.00 | 0.00 | 0.00 | 0.00 | 0.19 | 1.49 | 1,398 | 0.137 | 0 | 0 |
|  |  | **Arousal ratings** | abuse | 0.05 | 0.04 | -0.03 | 0.12 | 0.09 | 1.17 | 1,398 | 0.240 | 0 | 0 |
|  |  |  | neglect | 0.04 | 0.03 | -0.03 | 0.10 | 0.08 | 1.09 | 1,398 | 0.275 | 0 | 0 |
|  |  |  | interaction | 0.00 | 0.00 | 0.00 | 0.00 | -0.14 | -1.13 | 1,398 | 0.258 | 0 | 0 |
|  |  | **Valence ratings** | abuse | 0.04 | 0.04 | -0.04 | 0.12 | 0.08 | 1.03 | 1,398 | 0.303 | 0 | 0 |
|  |  |  | neglect | 0.03 | 0.03 | -0.04 | 0.10 | 0.07 | 0.88 | 1,398 | 0.382 | 0 | 0 |
|  |  |  | interaction | 0.00 | 0.00 | 0.00 | 0.00 | -0.13 | -1.02 | 1,398 | 0.309 | 0 | 0 |
|  |  | **Contingency ratings** | abuse | 0.81 | 0.58 | -0.33 | 1.95 | 0.11 | 1.39 | 1,398 | 0.165 | 0 | 0 |
|  |  |  | neglect | 0.18 | 0.49 | -0.78 | 1.14 | 0.03 | 0.37 | 1,398 | 0.712 | 0 | 0 |
|  |  |  | interaction | -0.03 | 0.02 | -0.08 | 0.01 | -0.18 | -1.43 | 1,398 | 0.154 | 0 | 0 |
|  | **GEN** | **SCR** | abuse | 0.00 | 0.00 | 0.00 | 0.01 | 0.05 | 0.64 | 1,398 | 0.525 | 0 | 0 |
|  |  |  | neglect | 0.00 | 0.00 | -0.01 | 0.00 | -0.11 | -1.42 | 1,398 | 0.156 | 0 | 0 |
|  |  |  | interaction | 0.00 | 0.00 | 0.00 | 0.00 | 0.05 | 0.38 | 1,398 | 0.705 | 0 | 0 |
|  |  | **Arousal ratings** | abuse | 0.03 | 0.04 | -0.06 | 0.12 | 0.05 | 0.63 | 1,398 | 0.527 | 0 | 0 |
|  |  |  | neglect | -0.01 | 0.04 | -0.08 | 0.06 | -0.01 | -0.20 | 1,398 | 0.838 | 0 | 0 |
|  |  |  | interaction | 0.00 | 0.00 | 0.00 | 0.00 | -0.04 | -0.28 | 1,398 | 0.782 | 0 | 0 |
|  |  | **Valence ratings** | abuse | 0.04 | 0.05 | -0.05 | 0.13 | 0.06 | 0.83 | 1,398 | 0.405 | 0 | 0 |
|  |  |  | neglect | -0.02 | 0.04 | -0.10 | 0.06 | -0.04 | -0.47 | 1,398 | 0.636 | 0 | 0 |
|  |  |  | interaction | 0.00 | 0.00 | 0.00 | 0.00 | -0.01 | -0.10 | 1,398 | 0.918 | 0 | 0 |
|  |  | **Contingency ratings** | abuse | 0.56 | 0.53 | -0.48 | 1.60 | 0.08 | 1.05 | 1,398 | 0.293 | 0 | 0 |
|  |  |  | neglect | -0.26 | 0.44 | -1.13 | 0.62 | -0.04 | -0.58 | 1,398 | 0.564 | 0 | 0 |
|  |  |  | interaction | -0.01 | 0.02 | -0.06 | 0.03 | -0.09 | -0.67 | 1,398 | 0.502 | 0 | 0 |
| **LDS** | **GEN** | **SCR** | abuse | 0.00 | 0.00 | 0.00 | 0.00 | -0.03 | -0.38 | 1,398 | 0.702 | 0 | 0 |
|  |  |  | neglect | 0.00 | 0.00 | 0.00 | 0.00 | -0.06 | -0.77 | 1,398 | 0.443 | 0 | 0 |
|  |  |  | interaction | 0.00 | 0.00 | 0.00 | 0.00 | 0.07 | 0.53 | 1,398 | 0.596 | 0 | 0 |
|  |  | **Arousal ratings** | abuse | -0.01 | 0.02 | -0.06 | 0.04 | -0.03 | -0.35 | 1,398 | 0.726 | 0 | 0 |
|  |  |  | neglect | -0.01 | 0.02 | -0.05 | 0.03 | -0.04 | -0.50 | 1,398 | 0.618 | 0 | 0 |
|  |  |  | interaction | 0.00 | 0.00 | 0.00 | 0.00 | 0.06 | 0.43 | 1,398 | 0.667 | 0 | 0 |
|  |  | **Valence ratings** | abuse | 0.00 | 0.02 | -0.05 | 0.05 | 0.00 | 0.02 | 1,398 | 0.982 | 0 | 0 |
|  |  |  | neglect | 0.01 | 0.02 | -0.03 | 0.05 | 0.04 | 0.51 | 1,398 | 0.612 | 0 | 0 |
|  |  |  | interaction | 0.00 | 0.00 | 0.00 | 0.00 | -0.04 | -0.34 | 1,398 | 0.738 | 0 | 0 |
|  |  | **Contingency ratings** | abuse | -0.04 | 0.30 | -0.63 | 0.55 | -0.01 | -0.14 | 1,398 | 0.891 | 0 | 0 |
|  |  |  | neglect | -0.16 | 0.25 | -0.66 | 0.33 | -0.05 | -0.64 | 1,398 | 0.523 | 0 | 0 |
|  |  |  | interaction | 0.01 | 0.01 | -0.02 | 0.03 | 0.07 | 0.52 | 1,398 | 0.603 | 0 | 0 |
| **General reactivity** | **ALL** | **SCR** | abuse | 0.00 | 0.00 | 0.00 | 0.01 | 0.03 | 0.42 | 1,398 | 0.676 | 0 | 0 |
|  |  |  | neglect | 0.00 | 0.00 | -0.01 | 0.00 | -0.11 | -1.42 | 1,398 | 0.156 | 0 | 0 |
|  |  |  | interaction | 0.00 | 0.00 | 0.00 | 0.00 | 0.04 | 0.28 | 1,398 | 0.778 | 0 | 0 |
|  |  | **Arousal ratings** | abuse | 0.04 | 0.02 | 0.00 | 0.09 | 0.15 | 1.94 | 1,398 | 0.053 | 0 | 0 |
|  |  |  | neglect | 0.01 | 0.02 | -0.02 | 0.05 | 0.06 | 0.74 | 1,398 | 0.459 | 0 | 0 |
|  |  |  | interaction | 0.00 | 0.00 | 0.00 | 0.00 | -0.19 | -1.50 | 1,398 | 0.133 | 0 | 0 |
|  |  | **Valence ratings** | abuse | 0.01 | 0.02 | -0.02 | 0.05 | 0.06 | 0.73 | 1,398 | 0.468 | 0 | 0 |
|  |  |  | neglect | -0.01 | 0.01 | -0.04 | 0.02 | -0.05 | -0.69 | 1,398 | 0.493 | 0 | 0 |
|  |  |  | interaction | 0.00 | 0.00 | 0.00 | 0.00 | -0.04 | -0.27 | 1,398 | 0.785 | 0 | 0 |
|  |  | **Contingency ratings** | abuse | -0.10 | 0.26 | -0.60 | 0.40 | -0.03 | -0.40 | 1,180 | 0.689 | 0 | 0 |
|  |  |  | neglect | -0.17 | 0.22 | -0.60 | 0.25 | -0.07 | -0.80 | 1,180 | 0.424 | 0 | 0 |
|  |  |  | interaction | 0.01 | 0.01 | -0.01 | 0.03 | 0.13 | 0.94 | 1,180 | 0.350 | 0 | 0 |
| Note. ACQ = acquisition training, GEN = generalization phase, LDS = linear deviation score. Bold numbers indicate significant results (p < 0.05). | | | | | | | | | | | | | |

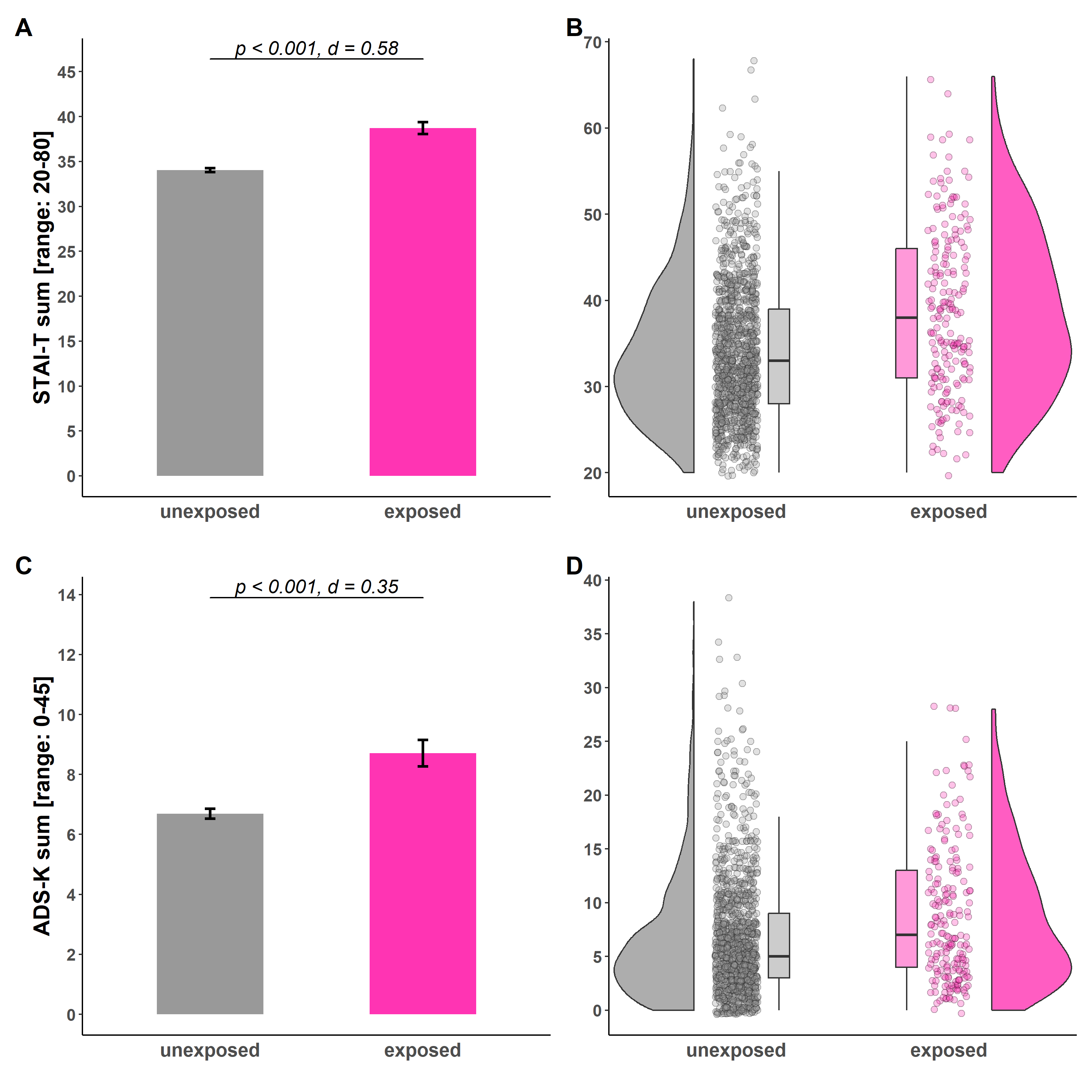
Supplementary Figure 6: Illustration of STAI-T (A-B) and ADS-K (C-D) sum scores for individuals unexposed (gray) and exposed (pink) to childhood adversity. Barplots (A and C) with error bars represent means and SEMs. with n_unexposed_ = 1199 and n_exposed_ = 203, respectively. The statistical parameters presented in A) and C) are derived from Welch's tests. The a priori significance level was set to α = 0.05. Distributions of the data are illustrated in the raincloud plots (B and D). Points next to the densities represent the sum scores of each participant. Boxes of boxplots represent the interquartile range (IQR) crossed by the median as a bold line, ends of whiskers represent the minimum/maximum value in the data within the range of 25th/75th percentiles ±1.5 IQR. STAI-T = Trait scale of the State-Trait Anxiety Inventory (Spielberger, 1983); ADS-K = Allgemeine Depressionsskala - Kurzform (short version of the Center for Epidemiological Studies-Depression Scale, CES-D; Allgemeine Depressions-Skala, ADS-K, Hautzinger & Bailer, 1993).

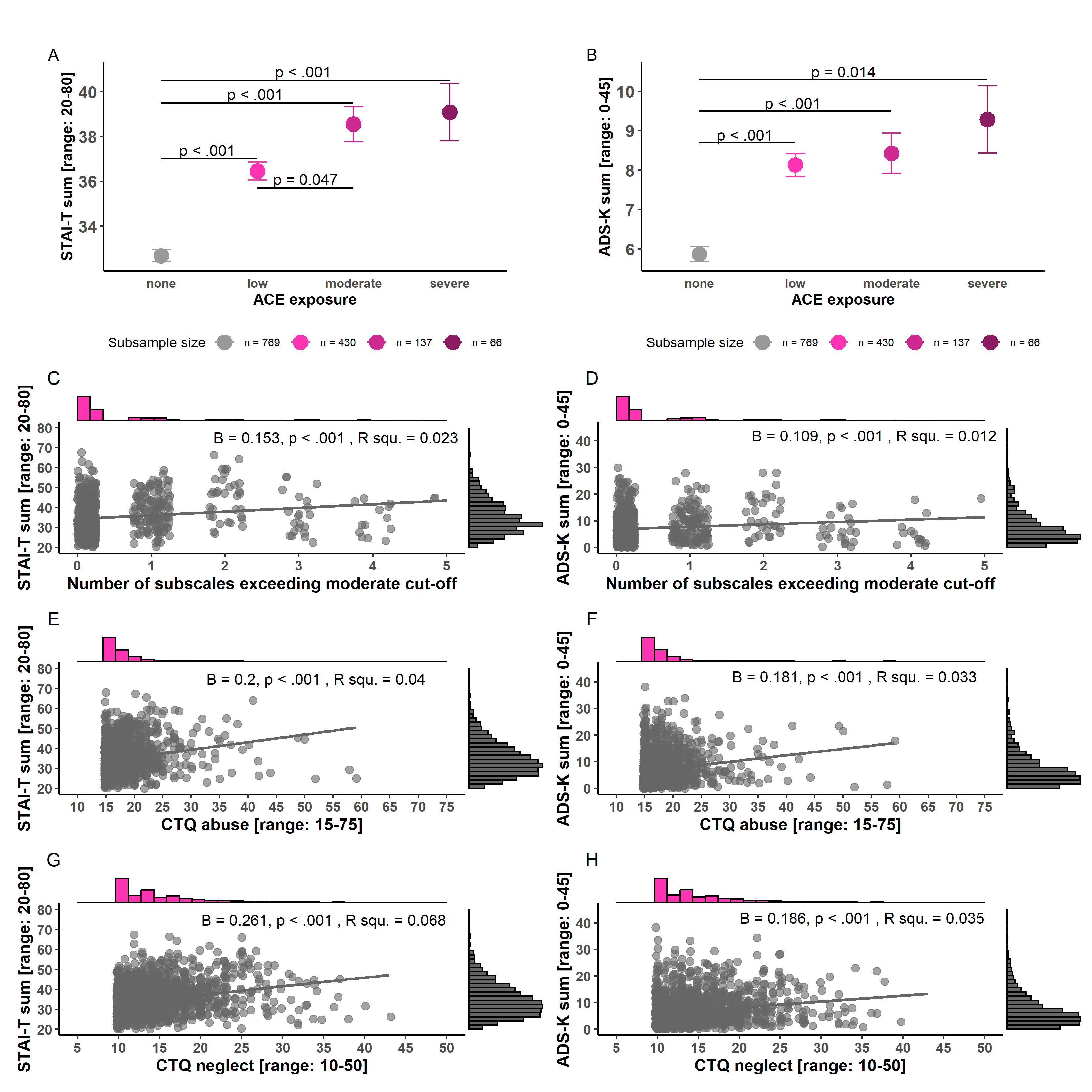

Supplementary Figure 7: Illustration of associations of STAI-T (A, C, E, G) and ADS-K (B, D, F, H) with childhood adversity operationalized as cumulative risk models (A - D) and the specificity model (E - H). STAI-T = Trait scale of the State-Trait Anxiety Inventory (Spielberger, 1983); ADS-K = Allgemeine Depressionsskala - Kurzform (short version of the Center for Epidemiological Studies-Depression Scale, CES-D; Allgemeine Depressions-Skala, ADS-K, Hautzinger & Bailer, 1993).

Supplementary Table 8: Pearson correlations of STAI-T and ADS-K with SCR

|  | **STAI-T** | **ADS-K** |
| --- | --- | --- |
| CS discrimination in SCR (ACQ) | r = -0.05 , p = 0.06 | **r = -0.057 , p = 0.033** |
| SCR to the CS+ (ACQ) | **r = -0.057 , p = 0.032** | **r = -0.057 , p = 0.032** |
| SCR to the CS- (ACQ) | r = -0.019 , p = 0.467 | r = -0.013 , p = 0.638 |
| CS discrimination in SCR (GEN) | r = 0.001 , p = 0.964 | r = -0.032 , p = 0.234 |
| LDS in SCR | r = -0.045 , p = 0.091 | r = -0.038 , p = 0.153 |
| Mean reactivity in SCR | r = -0.019 , p = 0.484 | r = -0.006 , p = 0.835 |
| Note. ACQ = acquisition training; GEN = generalization phase; STAI-T = Trait scale of the State-Trait Anxiety Inventory (Spielberger, 1983); ADS-K = Allgemeine Depressionsskala - Kurzform (short version of the Center for Epidemiological Studies-Depression Scale, CES-D; Allgemeine Depressions-Skala, ADS-K, Hautzinger & Bailer, 1993); LDS = linear deviation score. Bold numbers indicate significant results (p < 0.05). | | |
